# The impact of sequencing depth on the inferred taxonomic composition and AMR gene content of metagenomic samples

**DOI:** 10.1101/593301

**Authors:** H. Soon Gweon, Liam P. Shaw, Jeremy Swann, Nicola De Maio, Manal AbuOun, Alasdair T. M. Hubbard, Mike J. Bowes, Mark J. Bailey, Tim E. A. Peto, Sarah J. Hoosdally, A. Sarah Walker, Robert P. Sebra, Derrick W. Crook, Muna Anjum, Daniel S. Read, Nicole Stoesser, on behalf of the REHAB consortium

**Affiliations:** Harborne Building, School of Biological Sciences, University of Reading, RG6 6AS, UK; Centre for Ecology & Hydrology, Wallingford, Oxfordshire, OX10 8BB, UK; Nuffield Department of Medicine, University of Oxford, Oxford, UK; Department of Bacteriology, Animal and Plant Health Agency, Addlestone, Surrey, KT15 3NB, UK; NIHR Health Protection Research Unit (HPRU) in Healthcare-associated Infections and Antimicrobial Resistance at University of Oxford in partnership with Public Health England, Oxford, UK; Department of Genetics and Genomics, Icahn School of Medicine at Mt Sinai, New York, NY, USA

**Keywords:** Antimicrobial resistance (AMR), One Health, metagenomics, *Enterobacteriaceae*

## Abstract

**Background:** Shotgun metagenomics is increasingly used to characterise microbial communities, particularly for the investigation of antimicrobial resistance (AMR) in different animal and environmental contexts. There are many different approaches for inferring the taxonomic composition and AMR gene content of complex community samples from shotgun metagenomic data, but there has been little work establishing the optimum sequencing depth, data processing and analysis methods for these samples. In this study we used shotgun metagenomics and sequencing of cultured isolates from the same samples to address these issues. We sampled three potential environmental AMR gene reservoirs (pig caeca, river sediment, effluent) and sequenced samples with shotgun metagenomics at high depth (∼200 million reads per sample). Alongside this, we cultured single-colony isolates of *Enterobacteriaceae* from the same samples and used hybrid sequencing (short- and long-reads) to create high-quality assemblies for comparison to the metagenomic data. To automate data processing, we developed an open-source software pipeline, ‘ResPipe’.

**Results:** Taxonomic profiling was much more stable to sequencing depth than AMR gene content. 1 million reads per sample was sufficient to achieve <1% dissimilarity to the full taxonomic composition. However, at least 80 million reads per sample were required to recover the full richness of different AMR gene families present in the sample, and additional allelic diversity of AMR genes was still being discovered in effluent at 200 million reads per sample. Normalising the number of reads mapping to AMR genes using gene length and an exogenous spike of *Thermus thermophilus* DNA substantially changed the estimated gene abundance distributions. While the majority of genomic content from cultured isolates from effluent was recoverable using shotgun metagenomics, this was not the case for pig caeca or river sediment.

**Conclusions:** Sequencing depth and profiling method can critically affect the profiling of polymicrobial animal and environmental samples with shotgun metagenomics. Both sequencing of cultured isolates and shotgun metagenomics can recover substantial diversity that is not identified using the other methods. Particular consideration is required when inferring AMR gene content or presence by mapping metagenomic reads to a database. ResPipe, the open-source software pipeline we have developed, is freely available (https://gitlab.com/hsgweon/ResPipe).

## BACKGROUND

Antimicrobial resistance (AMR) is a significant global health threat (1,2) and understanding the evolution, emergence and transmission of AMR genes requires a ‘One Health’ approach considering human, animal and environmental reservoirs (3). Methods for profiling species and AMR gene content in samples from these niches can be broadly categorised as either culture-dependent or culture-independent. Culture-dependent methods have the advantage of isolating individual strains for detailed analysis, but hugely underestimate species and AMR gene diversity. Culture-independent methods typically involve shotgun metagenomics, in which the DNA of a complete microbial community in a sample is extracted and sequenced, and the sequencing reads are used to estimate AMR gene and/or species distributions. The advantage of shotgun metagenomics is its relative lack of bias, but it tends to be less sensitive than targeted, culture-based or molecular approaches identifying specific drug-resistant isolates or AMR genes of interest (4–6).

Problems in characterising the epidemiology of AMR are exemplified by the *Enterobacteriaceae* family of bacteria. This family contains over 80 genera, and includes many common human and animal pathogens, such as *Escherichia coli*, that can also asymptomatically colonise human and animal gastrointestinal tracts, and are also found in environmental reservoirs (7). The genetic diversity of some *Enterobacteriaceae* species is remarkable: in *E. coli*, it has been estimated that only ∼10% of the 18,000 orthologous gene families found in the pangenome are present in all strains (8). AMR in *Enterobacteriaceae* is mediated by >70 resistance gene families, and >2,000 known resistance gene variants have been catalogued (9,10). In addition to mutational resistance, AMR genes are also commonly shared both within and between species on mobile genetic elements such as insertion sequences, transposons and plasmids. Individuals have been shown to harbour multiple diverse AMR gene variants, strains and species of *Enterobacteriaceae* in their gastrointestinal tract (11,12), highlighting that single-colony subcultures do not recover the true AMR reservoir even within a small subsection of a microbial community.

Attempting to near-completely classify AMR gene and species diversity by any culture-based approach for raw faeces, effluent, and river sediment is therefore unlikely to be feasible; hence, the use of shotgun metagenomics to achieve this aim. However, the replicability of metagenomic surveys and the sequencing depth (reads per sample) required to analyse these sample types has not yet been explored in detail (13,14).

Motivated by the need to analyze large numbers of these samples in the REHAB study (http://modmedmicro.nsms.ox.ac.uk/rehab/), here we carried out a pilot study (Figure 1) to investigate: (i) the replicability of sequencing outputs using common DNA extraction and sequencing methods; and the impact of (ii) widely used taxonomic and AMR gene profiling approaches; (iii) sequencing depth on taxonomic and AMR gene profiles; and (iv) sequencing depth on the recoverability of genetic contents from isolates identified in the same samples using culture-based approaches.

**Figure 1.**
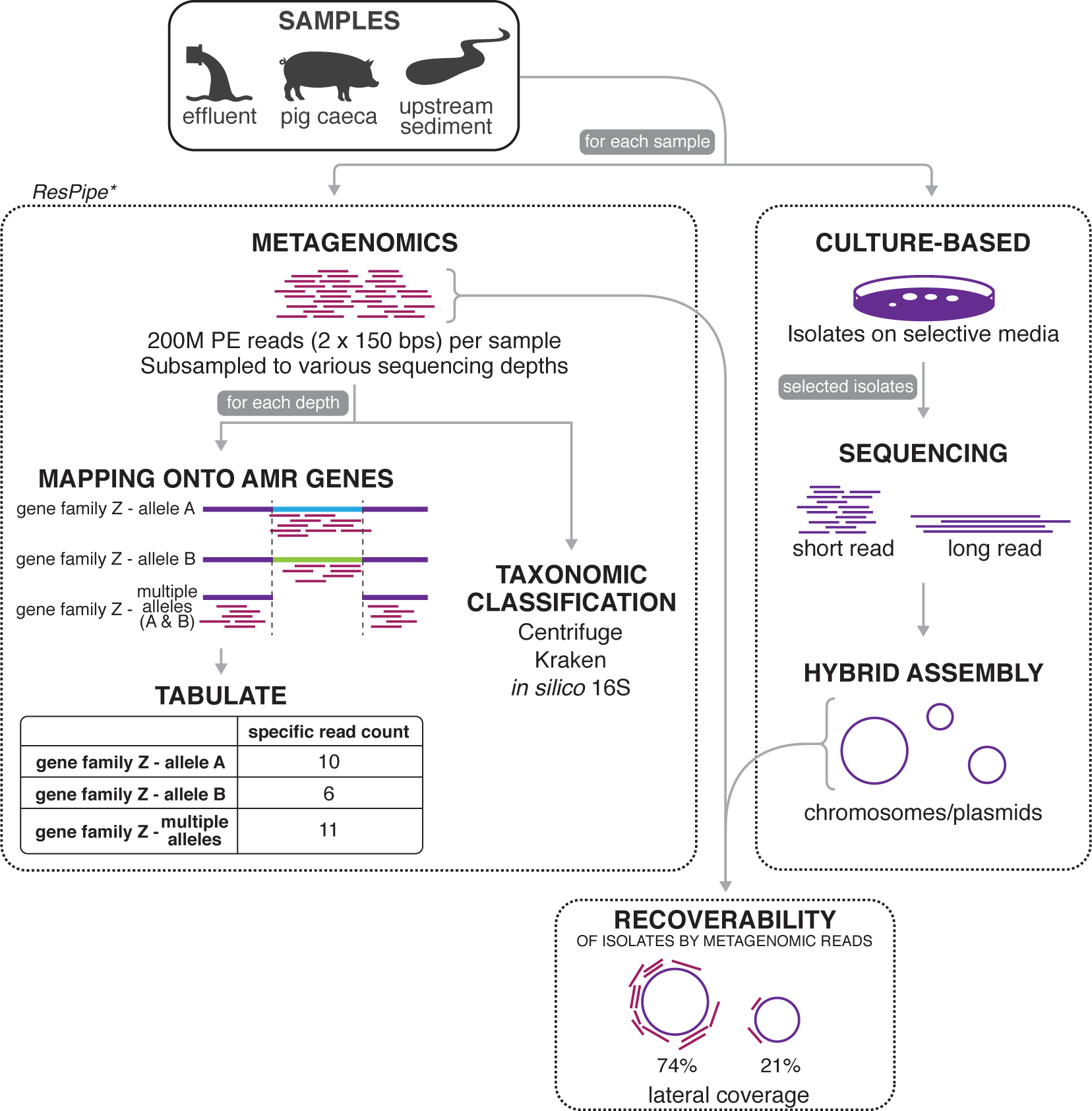
Schematic overview of the study. For each sample, we used both a metagenomics and culture-based approach. We developed a software pipeline (‘ResPipe’) for the metagenomic data. For more details on each step of the workflow, see Methods.

## RESULTS

### Impact of sequencing depth on AMR profiles

Metagenomic sequencing produced approximately 200 million metagenomic 150bp paired-end reads per sample i.e. over 56 gigabases per sample (Supplementary Table 1), of which <0.05% of reads mapped with 100% identity to a known AMR-related sequence (see next section). The number of reads mapping to AMR gene families was largest in pig caeca (88,816 reads) and effluent (77,044 reads). Upstream sediment did not have enough AMR-related reads for further analysis (49 reads).

The effluent sample had the highest total richness of both AMR gene families and AMR allelic variants (Figure 2). Sequencing depth significantly affected the ability to evaluate richness of AMR gene families in effluent and pig caeca, which represent highly diverse microbial environments. The number of AMR gene families observed in effluent and pig caeca stabilized (see Methods: ‘Rarefaction curves’) at a sequencing depth of ∼80 million reads per sample (depth required to achieve 95% of estimated total richness, *d*_0.95_: 72-127 million reads per sample). For AMR allelic variants in effluent, the richness did not appear to have plateaued even at a sequencing depth of 200 million reads per sample, suggesting the full allelic diversity was not captured (*d*_0.95_: 193 million reads per sample).

**Figure 2.**
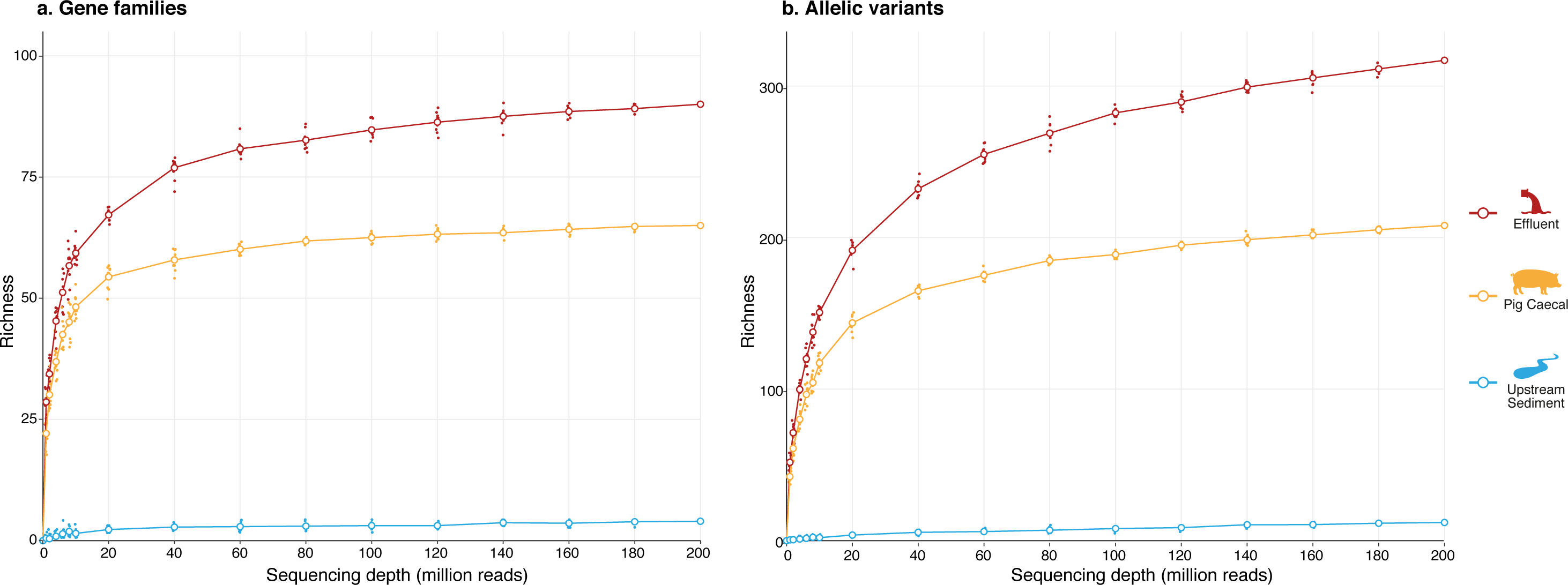
Rarefaction curve at various sequencing depths for (a) AMR gene families, and (b) AMR gene allelic variants. Colours indicate sample type. For each sampling depth, sequences were randomly subsampled 10 times, with each point representing a different subsampling. Lines connect the means (large circles) of these points for each sample type.

### Specific mapping to AMR genes and allelic variants

We exploited the hierarchical structure of the Comprehensive Antimicrobial Resistance Database (CARD) to assign reads to their respective AMR gene families and AMR allelic variants using a specific read mapping strategy i.e. to count only reads which mapped to a unique region of an allele or a gene family. In order to place a lower bound on the AMR diversity present, we adopted a stringent approach which counted only alignments with 100% sequence identity to CARD sequences. The resulting AMR gene family profiles differed significantly between the samples (Figure 3). The most abundant AMR gene families in effluent and pig caeca were “23S rRNA with mutations conferring resistance to macrolide” and “tetracycline-resistant ribosomal protection protein”, respectively. There were 10,631 and 733 reads assigned to a “multiple gene family” category in the effluent and pig caeca, respectively. These represent reads that were mapped across multiple AMR gene families and therefore could not be uniquely assigned to any single family.

**Figure 3.**
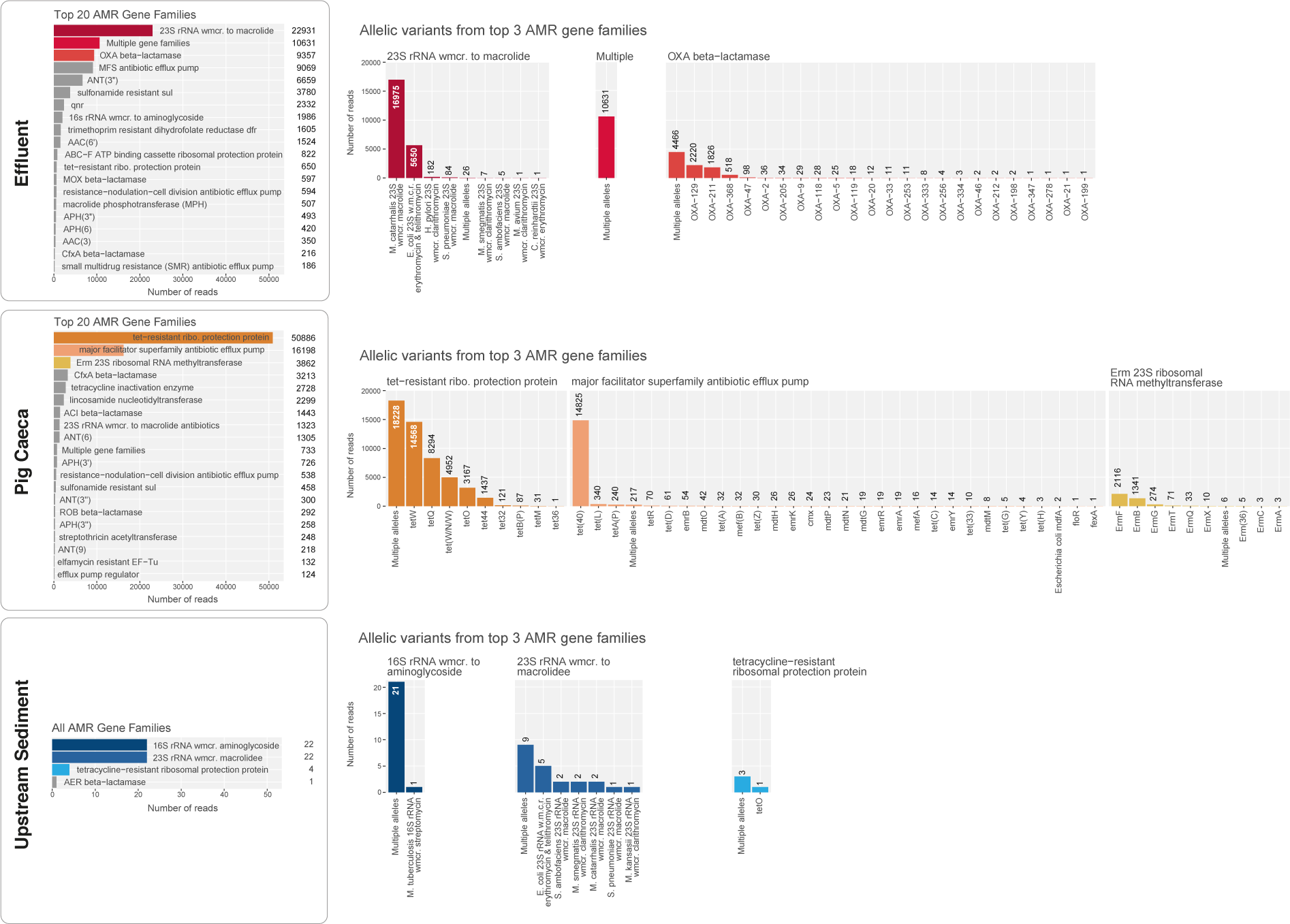
The most common AMR gene families and gene allelic variants in each sample. Left panel: the top 20 AMR gene families from effluent, pig caeca and upstream sediment by number of reads (top to bottom), with the top three most abundant highlighted in colour (hue indicates sample type) for comparison with the right-hand panel. Right panel: the most abundant AMR gene allelic variants within these top three most abundant gene families (left to right), sorted by abundance. For more information on the definitions of ‘AMR gene family’ and ‘allelic variant’, see Methods: ‘AMR gene profiling’.

Reads that mapped to one specific AMR gene family but onto multiple allelic variants (i.e. could not be assigned to one specific allele) were classified as “multiple alleles”. There was evidence of high allelic diversity, including among clinically relevant AMR gene families. For example, 47.7% of the reads mapped to the “OXA beta-lactamase” family could not be assigned to a specific allele (4,466 out of 9,357 reads; third-most abundant gene family by reads). Similarly, the most abundant gene family by reads in pig caeca was “tetracycline-resistant ribosomal protection protein”, and 35.8% of the reads that mapped within this family could not be assigned to a specific allele (18,228 out of the 50,886 reads).

### Impact of normalisation strategies on AMR allelic variant abundances

Normalising by gene length (see Methods: ‘Normalisation of gene counts’) had a profound effect on the distributions and the ranking order of AMR allelic variants in general (Figure 4). Further normalisation by *T. thermophilus* reads did not affect the per sample distributions of AMR allelic variants, but it allowed more accurate comparison between samples by estimating absolute abundance of any given variant in the sample. The number of reads that mapped to *T. thermophilus* were similar between three samples, and this meant that the changes were small (i.e. a slight relative increase in the effluent compared to the pig caeca sample). While most of the alleles had lateral coverages between 90% and 100% in effluent and pig caeca samples (Figure 3, right panels), “*Moraxella catarrhalis* 23S rRNA with mutation conferring resistance to macrolide antibiotics” had lateral coverage of 29% despite being one of the most abundant alleles in the effluent.

**Figure 4.**
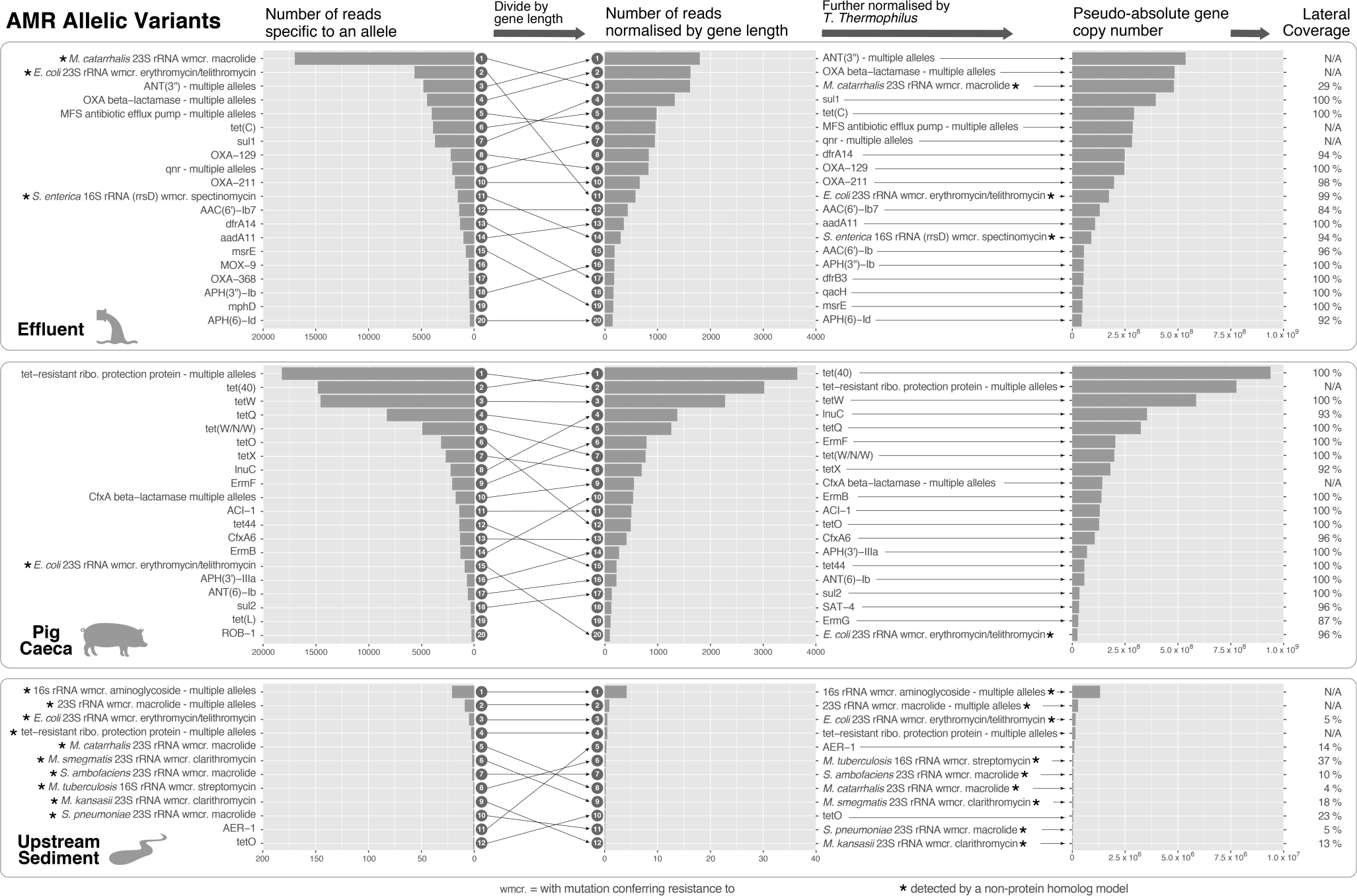
The effect of normalization on the most common AMR gene allelic variants from each sample. Shown are the top 20 AMR gene allelic variants from each sample (effluent, pig caeca and upstream sediment), and the effect of different normalisations (left: raw count, middle: normalisation by gene length, right: further normalisation by *Thermus thermophilus* count). Arrows show the changing rank of each variant with normalisation. Note that a different x-axis is used for upstream sediment in all three panels. Asterisks denote AMR allelic variants that do not have a “protein homolog” detection model in CARD (see Methods: ‘AMR gene profiling’).

### Impact of different assignment methods on taxonomic composition

While Centrifuge overall classified more reads than Kraken, both methods showed a similar trend of effluent having a greater proportion of reads classified as bacterial compared to upstream sediment, which had more than pig caeca (Figure 5a). Apart from Centrifuge classifying noticeably more Eukaryota and Viruses (0.7% and 0.05% respectively) than Kraken (0.09% and 0.01% respectively), a large proportion of reads from both methods were unclassified (70.0% and 83.3% for Centrifuge and Kraken respectively). The proportions of recoverable bacterial 16S rRNA fragments were low for all samples (0.16%, 0.23% and 0.04% for effluent, pig caeca and upstream sediment samples respectively), highlighting that shotgun metagenomics is an extremely inefficient method for obtaining 16S rRNA gene sequences.

**Figure 5.**
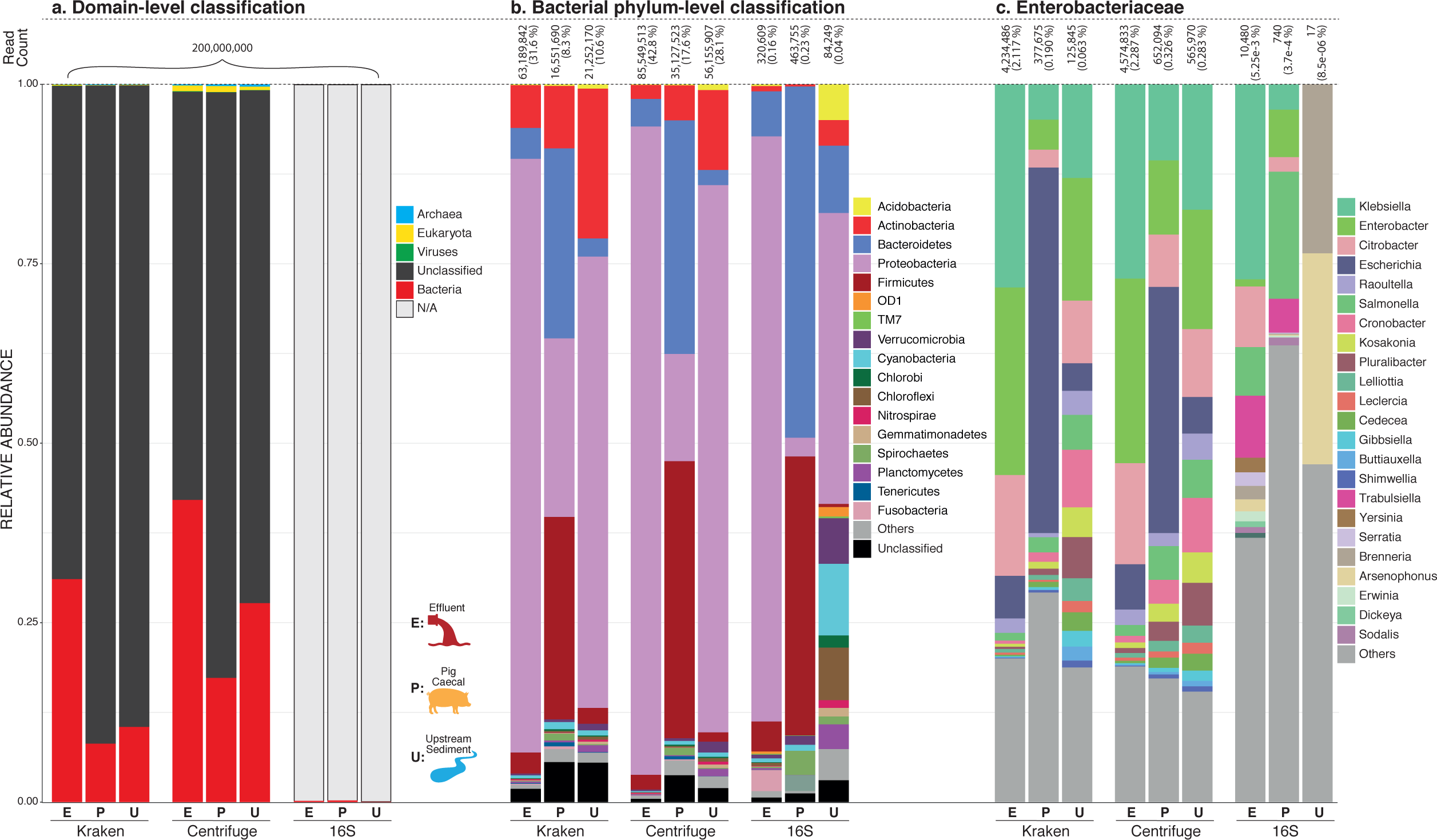
Taxonomic classification of metagenomes by method. Resulting taxonomic composition of effluent (E), pig caeca (P) and upstream sediment (U) metagenomes using Kraken, Centrifuge and classification by *in silico* 16S rRNA extraction (16S). (a) Domain-level classification. (b) Relative abundance of bacterial phyla (c) Relative abundance of *Enterobacteriaceae*.

The bacteria phylum-level classication (Figure 5b) showed structural differences among all three classification methods. The overall community structure and composition were more similar between Kraken and Centrifuge than the ‘*in silico* 16S’ approach (see Methods: ‘Taxonomic profiling’). This was particularly apparent in the upstream sediment, where using ‘*in silico* 16S’ produced distinctively different community profiles from the other methods. Kraken and Centrifuge classified between 377,675 to over 4 million reads as *Enterobacteriaceae*. Again, overall composition was similar between these two methods but showed some granularity in structure for pig caeca e.g. the relative abundances of *Escherichia* were 34.3% and 50.9%, and for *Klebsiella* 10.6% and 4.9%, for Centrifuge and Kraken respectively.

### Impact of sequencing depth on genus-level richess and taxonomic profiles

Kraken and Centrifuge taxonomic profiles were highly stable to sequencing depth within samples. Comparing different sequencing depths within samples using Bray-Curtis dissimilarity showed that the relative taxonomic composition was highly robust to sequencing depth, with 1 million reads per sample already sufficient for <1% dissimilarity to the composition inferred from 200 million reads per sample (Supplementary Figure 1). This was true at both the genus and species level, even though all classification methods are known to have less precision and sensitivity at the species level (15,16). Intriguingly, the genus-level richness rapidly reached a plateau for all samples at ∼1 million reads per sample (Figure 6a and Figure 6b), suggesting a database artifact (see ‘Discussion’).

**Figure 6.**
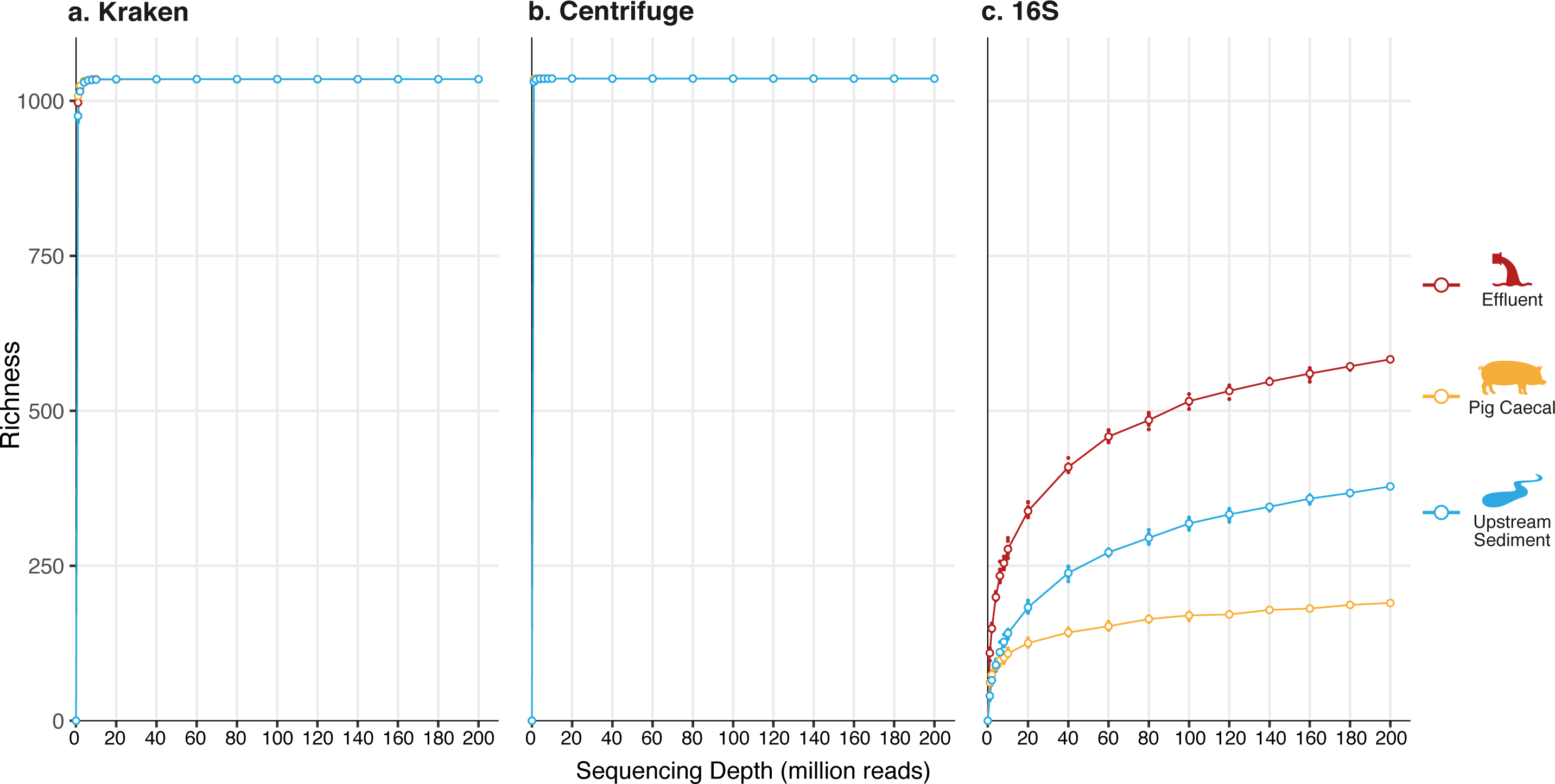
Impact of sequencing depth on genus-level richness. Three methods are shown: (a) Kraken, (b) Centrifuge and (c) *in silico* 16S rRNA extraction.

### Recovery of known genomic structures from cultured isolates using metagenomes

In order to assess how well shotgun metagenomics could recapitulate culture-dependent diversity, we cultured seven *Enterobacteriaeceae* isolates (four from effluent, two from pig caeca, one from upstream sediment; Table 1), then performed hybrid assembly (Supplementary Table 2). We then assembled near-complete genomes and mapped metagenomic reads back to these genomes (see Methods: ‘Mapping of metagenomic sequences onto isolates’; Supplementary Table 3). 26/28 contigs from effluent isolates rapidly achieved 100% lateral coverage at 1X using metagenomic reads at 80-100 million reads per sample (Figure 7a), with the two other contigs having almost-complete coverage at 200 million reads (98.7% and 99.8% respectively). Pig caeca isolates showed lower but fairly comprehensive lateral coverage of at least 75% for chromosomes at 200 million reads (Figure 7b), but only one contig (P1-5, shown in yellow) reached complete lateral coverage. The single chromosomal contig recovered from the upstream sediment isolate only had 0.2% of its bases covered at 200 million reads per sample, reflecting its scarcity in the metagenome (Figure 7c, Supplementary Table 3).

**Figure 7.**
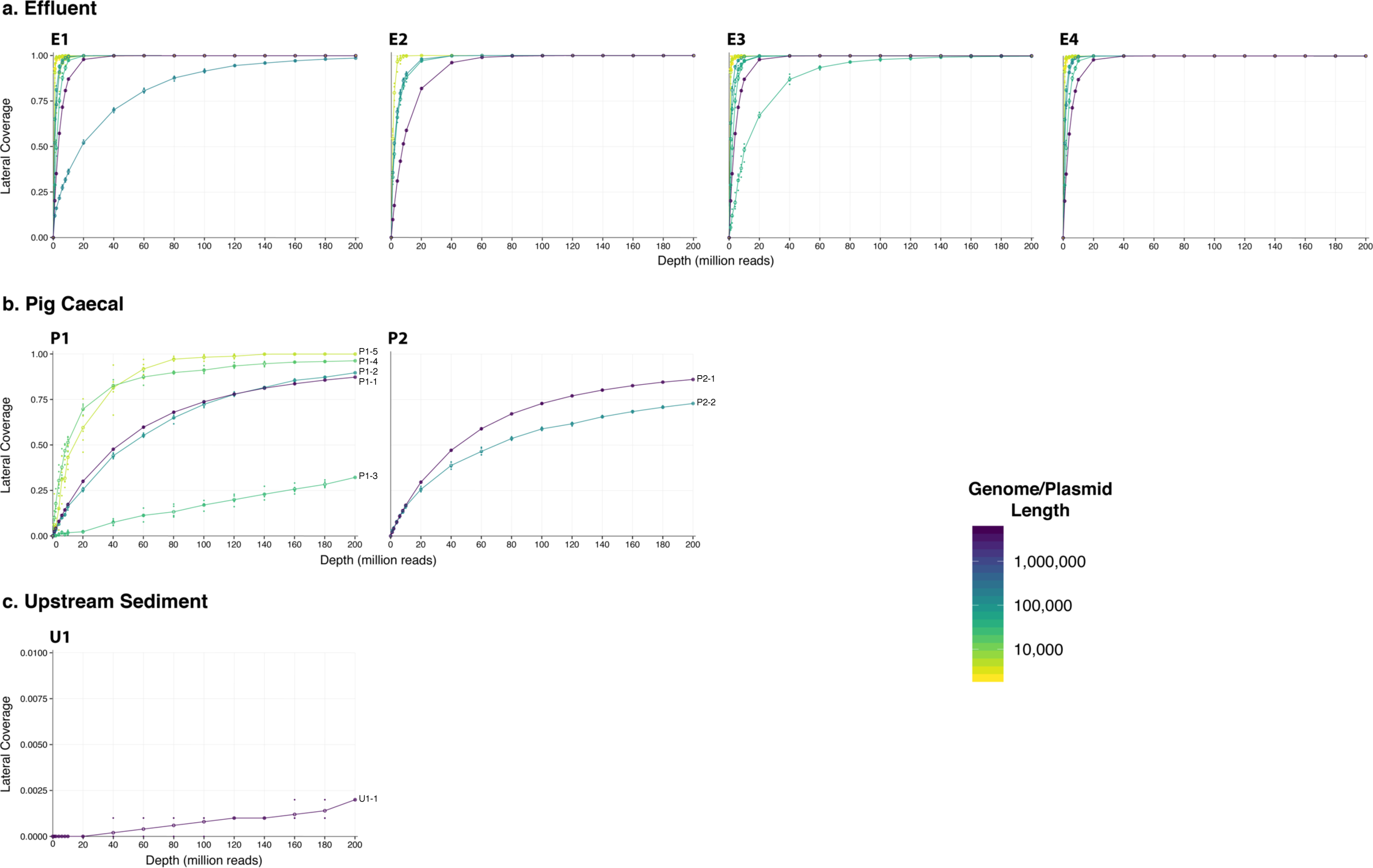
Metagenomic read coverage of assembled genetic structures from isolates cultured from each sample. (a) Effluent isolates: E1-E4, (b) Pig caeca isolates: P1-P2, (c) Upstream sediment isolate: U1. Genetic structures are coloured by size. Note the different y-axis scale for the upstream sediment sample.

**Table 1.** Details of cultured isolates and assembled genomes. For more details on isolate sequencing, see Supplementary Table 4.

| Sample | Isolate number | Species (MALDI-ToF) | Genome size (bp) | Number of contigs (number circularized) |
| --- | --- | --- | --- | --- |
| Effluent | 1 | *Citrobacter braakii* | 5,213,846 | 4 (4) |
|  | 2 | *Enterobacter kobei* | 5,590,302 | 9 (7) |
|  | 3 | *Enterobacter cloacae* | 5,465,276 | 7 (5) |
|  | 4 | *Enterobacter cloacae* | 5,393,186 | 8 (4) |
| Pig caeca | 1 | *E. coli*/S. *dysenteriae* | 4,898,477 | 5 (5) |
|  | 2 | *E. coli*/S. *dysenteriae* | 4,967,077 | 2 (2) |
| Upstream sediment | 1 | *Citrobacter gillenii* | 4,839,493 | 1 (1) |

## DISCUSSION

To our knowledge, our study is the first to have simultaneously investigated effluent, animal caecal and environmental metagenomics with deep sequencing of 200 million 150 bp paired-end reads per sample (∼60 gigabases per sample). Previous studies have used from 10 million to 70 million reads per sample (approximate bases per sample: 3Gb (17), 4 Gb (19), 7 Gb (6), 12 Gb (18)), often with shorter reads. We have demonstrated the significant effect of sequencing depth on taxonomic and AMR gene content profiling, and the ability to recover genomic content (obtained via single-colony culture of isolates from the sample) from metagenomics. In brief, we find that while accurately capturing broad-scale taxonomic composition requires relatively low sequencing depth, this is emphatically not the case for AMR gene diversity. This has critical importance for the many studies that seek to characterise animal and environmental reservoirs of AMR, and for the contextualisation of findings reported in previous metagenomics studies.

Deep metagenomic sequencing has been investigated more thoroughly in the context of the human microbiome. Hillmann et al. (2018) recently reported ultradeep metagenomics (2.5 billion reads) on two human stool samples, concluding that as few as 0.5 million reads per sample could recover broad-scale taxonomic changes and species profiles at >0.05% relative abundance (14). In line with this, we find that 1 million reads per sample is already sufficient to accurately obtain taxonomic composition (at <1% dissimilarity to the ‘true’ composition at 200 million reads). However, even 200 million reads per sample is not enough to obtain the complete diversity of AMR genes in effluent. This is potentially concerning because environmental metagenomics studies often use sequencing depths of as little as ∼10 million reads per sample (∼3.6Gb). For pig caeca samples, 80 million reads per sample appears to be adequate for sampling all AMR gene families represented in CARD, but still not adequate for exhausting AMR allelic variants. Notably, we adopted the stringent criterion of a perfect (i.e. 100%) match to assign any given read to a reference AMR sequence. This strategy obviously reduces the risk of false positives, while increasing false negatives. Therefore, our results represent a conservative lower bound on the AMR diversity present in the samples we analysed.

An additional challenge of metagenomics analysis in the context of AMR is choosing a consistent strategy for ‘counting’ AMR genes, whether in terms of their presence or relative abundance, from mapped reads. It remains unclear what the best approach is for this problem. One option is to count all the reads which map to a reference gene; however, this means that reads are potentially counted multiple times when the reference gene shares homology with other genes in the database, or that counts may be underestimated if reads are randomly assigned to best reference matches. In addition, reads which map to a wildtype, non-resistant sequence may also be inadvertently and inappropriately counted. Another option is to use only reads which map to regions of a gene that are unique and not shared with other genes in the database (e.g. as in ShortBRED (20)). This is a more conservative approach, but may be inherently biased against closely-related genes in the database. For example, CARD contains 14 sequences for *bla*_NDM_ genes, which differ at less than 2% of their positions, so each gene individually has very few specific regions. Exploiting knowledge of the often complex genetic variation within AMR gene families is necessary to avoid erroneous conclusions regarding presence/absence. Inferred abundances of particular AMR genes are likely frequently contingent not only on mapping and counting strategies, but also on the particular genetic features of the AMR genes catalogued in the chosen reference database. Interpreting and comparing results across studies utilising different methods therefore becomes difficult.

Once the type of count data to be considered (in terms of number of reads mapping to a gene) has been chosen, a normalisation strategy is required to compare across genes and samples. We found that normalising by gene length changed the inferred abundance distributions of AMR genes across all the sample types studied, again with important implications for those studies that have not undertaken this kind of normalisation. We have also outlined a protocol to obtain a pseudo-absolute gene copy number of specific regions of AMR genes by normalising by both gene length and an exogenous spike of *T. thermophilus*. While we do not claim that this accurately reflects the true abundance of individual genes, we believe it is useful for comparisons across samples within a study. In our study we took great care to ensure standardised DNA extraction and had small batches of samples; probably as a result, we obtained similar proportions of sequences of *T. thermophilus* for all samples (range: 0.067-0.082%), but this may not always be the case. Appropriate normalisation using exogenous DNA spikes to account for some of the extraction biases could have potentially dramatic effects on results and their interpretation.

As well as examining normalised abundances, the lateral coverage of a gene is also an important metric to decide whether a certain allele is likely present in the sample. In effluent, the most abundant gene by specific read count was “*Moraxella catarrhalis* 23S rRNA with mutation conferring resistance to macrolide antibiotics”. However, the gene only had 29% lateral coverage, and this result should therefore be interpreted cautiously. In fact, the high specific read count is probably because CARD only includes one *Moraxella* rRNA gene with an AMR mutation compared to twenty *Escherichia* rRNA genes; the lateral coverage suggests that the AMR allele is not in fact present. This underlines the importance of considering multiple metrics simultaneously.

Both taxonomic and AMR gene profiling outputs are clearly dependent on the species and AMR databases used as references. It should be additionally noted that for AMR gene profiling, some genes are variants of a wildtype which may differ by as little as a single SNP. Because short-read metagenomics typically surveys ≤150 bp fragments, even specific read counts can in fact plausibly be wildtypes rather than particular resistance variants. This can be overcome by adopting our stringent approach which requires an exact match (i.e. at 100%) to call a given variant in the database; although obviously this increases the rate of false negatives, we have shown that this strategy appears successful given adequate sequencing depth.

We found a reasonable consistency between taxonomic classification methods, but there were differences between Kraken and Centrifuge, and undoubtedly there would have been differences with other methods, had we tested them. This is a previously recognised issue (e.g. as in (21)) and has no single solution; methods are optimised for different purposes and perform differently depending on the combination of sample type, sequencing method, and reference database used. As the field changes so rapidly and newer methods become available, we strongly recommend that researchers with shotgun metagenomic data review excellent benchmarking efforts such as CAMI (21) and LEMMI (22) and assess the tools using a particular quantitative metric rather than making a (perhaps arbitrary) choice for their analysis. Investigating the robustness of conclusions to choice of method is also a recommended step (23,24).

Remarkably, there were no ‘unique genera’ at high sequencing depth: reads assigned to all genera were present in all three sample types at high depth. We believe this is an artifact due to the limited number of genomes available in the species database used for the assignment methods. The RefSeq database contains complete genomes for 11,443 strains, but these represent only 1,065 genera. Our samples almost exhausted the entire genus space: the number of genera that were classified by Centrifuge was 1,036, and this number was the same for the effluent, pig caeca and upstream sediment samples, i.e. all three samples had the same number of total unique genera observed at 200 million reads depth. This was the same with Kraken, which classified 1,035 genera in total and there was no difference in richness between the three samples. This highlights the importance of using diversity measures which take into account the relative abundance of taxa rather than just their presence or absence.

We also found that a large number of reads (>50%) were unclassified by either Kraken or Centrifuge. The absence of organisms such as fungi from our reference database could have played a role in this, but other studies of effluent have also found that between 42% and 68% of short metagenomic reads cannot be assigned to any reference sequence (25–27). Our focus was on using the best available tools to assess the bacterial composition of samples; understanding what this unassigned microbial ‘dark matter’ represents was beyond the scope of this study, but would be valuable future work.

Our analyses confirm that using culture-based methods offered complementary and additional information to shotgun metagenomics. By mapping metagenomic reads back to high-quality hybrid assemblies obtained via culture, we found that the majority of isolates from effluent were recoverable by metagenomics at sequencing depths. However, the majority of isolates from pig caeca were not recovered. These results exemplify the need for exploring both methods in analysing AMR genes and microbial communities, as both show different perspectives on the AMR profiles and strains present in a given sample.

## CONCLUSIONS

In summary, we have used a combination of deep metagenomic sequencing, hybrid assembly of cultured isolates, and taxonomic and AMR gene profiling methods to perform a detailed exploration of methodological approaches to characterise animal and environmental metagenomic samples. Sequencing depth critically affects the inferred AMR gene content and taxonomic diversity of complex, polymicrobial samples, and even 200 million reads per sample was insufficient to capture total AMR allelic diversity in effluent. Choice of taxonomic profiler can result in significant differences in inferred species composition.

The open-source software pipeline we have developed is freely available as ‘ResPipe’. As well as packaging existing tools, ResPipe provides detailed information on various metrics that are useful for assessing AMR gene abundances, including: a novel normalisation technique for read counts, specific mapping counts, and lateral coverage, all of which can provide different but important insights. There is undoubtedly vast diversity present in microbial communities. Establishing best practices and pipelines for analysing this diversity with shotgun metagenomics is crucial to appropriately assess AMR in environmental, animal and human faecal samples.

## METHODS

### Sample types and settings

We sampled three distinct potential AMR reservoirs, namely: (i) pooled pig caecal contents from 10 pigs from a breeder farm in Yorkshire and the Humber (denoted as “pig caeca”); (ii) river sediment 100m upstream of a sewage treatment works (STW) at Cholsey STW, Cholsey, Oxfordshire (“upstream sediment”); and (iii) treated sewage effluent emitted from Cholsey STW (“effluent”). Cholsey STW is a plant that serves a population equivalent of ∼21,000 with a consented flow of 3200 m^3^/day; processes include primary settlement tanks, followed by biological disc filters and humus tanks, and subsequently disc filtration. These sample types were chosen to represent a spectrum of predicted diversity of microbial communities (i.e. high to low: effluent, pig caeca, upstream sediment).

The pooled pig caeca had been collected as part of a separate study surveying the presence of AMR genes in *E. coli* in pigs from 56 farms across the UK (28). In brief, caecal contents were sampled from 10 randomly selected healthy finishing pigs from each of the farms at 12 different abattoirs (March 2014 - October 2015), and suspended in 22.5 mL of PBS (processing within 24hrs of collection). Aliquots of 100 µL were frozen at −80 °C. This study used an aliquot of pooled pig caeca selected randomly from this collection.

For effluent and upstream sediment samples, sterile Whirl-pack™ bags were attached to extendable sampling arms and placed into flow at the relevant site. Samples in the bags were stirred with sterile spoons, and 5 mLs added to a sterile 50 mL centrifuge tube. This process was repeated five times to create a composite sample of approximately 25 mL. Samples were stored in a cool box at 4 °C for transport and processed within 24 hrs.

### Metagenomic DNA extractions and *Thermus* spike-in

Metagenomic extractions on all samples were performed using the MoBio PowerSoil® DNA Isolation Kit (Qiagen, Venlo, Netherlands), as per the manufacturer’s protocol, and including a beadbeating step of two 40 second cycles at 6 m/s in lysing matrix E. 12.5 ng of naked *Thermus thermophilus* DNA (reference strain HB27, Collection number ATCC BAA-163, ordered from DSMZ, Germany) was added to each sample in the PowerBead tube at the start of the experiment, prior to the addition of Solution C1 of the DNA Isolation Kit. The rationale for this was to enable subsequent normalisation to the number of *T. thermophilus* genomes sequenced to adjust for varying amounts of sample input, and extraction bias (33) (see ‘Normalisation of gene counts’, below).

### Metagenomic sequencing

Pooled libraries of all DNA extracts were sequenced across four lanes of an Illumina HiSeq 4000 platform, generating a median of 102,787,432 150 bp paired-end reads (30.8 Gb) of data per extract. For the samples extracted in replicate, we therefore had a median of 202,579,676 paired-end reads (60.7 Gb) of data available for evaluation and sub-sampling analyses (Supplementary Table 1). To confirm replicability of our extraction method on the same sample, duplicate extractions of all three samples were performed. To test replicability of sequencing, pooled libraries derived from extracts were each sequenced across four sequencing lanes. The sequences were pooled into each sample resulting in 202,579,676, 215,047,930 and 198,865,221 reads for the effluent, pig caeca and upstream sediment respectively. The effluent and pig caeca samples were both randomly subsampled down to 200 million reads per sample for downstream analysis.

Analysis of both AMR gene profiles and taxonomic profiles for the same extract pooled across multiple sequencing lanes (HiSeq) were highly reproducible, with little evidence of differences across lanes, although there was a significant difference between replicates of AMR gene profiles from pooled pig caeca (*p*=0.03), and replicates of taxonomic profiles for upstream sediment (*p*=0.03) (Supplementary Table 4).

### Sequencing depth subsampling and quality filtering

In order to simulate the effect of sequencing at different depths, each set of pooled reads from the three samples was repeatedly subsampled (*n*=10) using VSEARCH (fastx_subsampling, (34)) into the following set of depth intervals: 1M, 2M, 4M, 6M, 7M, 8M, 9M, 10M, 20M, 40M, 60M, 80M, 100M, 120M, 140M, 160M and 180M. Low-quality portions of all reads were trimmed using TrimGalore (v.0.4.4_dev, (35)). Specifically, we used a length cut-off of 75 bp and average Phred score ≥25, and the first 13 bp of Illumina standard adapters (AGATCGGAAGAGC) for adapter trimming.

### Taxonomic profiling

For profiling the abundance of bacterial species, the reads were classified with Kraken (v.1.1, default settings; (16)) and Centrifuge (v.1.0.4, default settings; (15)), which were chosen based on recency and reported frequency of use in the literature. RefSeq sequences (v.91; (36)) at a “Complete genome” assembly level for bacteria (11,443 strains), archaea (275 strains), viral (7,855 strains) and human were downloaded from the NCBI repositories and used to build two sets of indexed databases for both Kraken and Centrifuge using respective scripts provided by each classifier. An ‘*in silico* 16S’ marker-gene based classification was performed by extracting 16S rRNA genes from the reads using METAXA2 (4) followed by taxonomic assignment with the naïve Bayesian RDP classifier (v2.10; (37)) with a minimum confidence of 0.5 against the GreenGenes database (v.13.5; (38)).

To validate the taxonomic profiling component of our pipeline, we analyzed ten previously simulated gut metagenomes (GI tract data from “2nd CAMI Toy Human Microbiome Project Dataset”, https://openstack.cebitec.uni-bielefeld.de:8080/swift/v1/CAMI_Gastrointestinal_tract) produced for benchmarking as part of CAMI (21). Comparing to the ground truth of the simulated composition, using either Centrifuge or Kraken recovered the major features of the taxonomic composition (Supplementary Figure 2a) with high correlation between simulated and inferred species abundances (Supplementary Figure 2b), although there were apparent discrepancies between methods which we did not investigate further.

### AMR gene profiling

The quality filtered reads were mapped with bbmapskimmer.sh (BBMap suite; (39)) with default settings against sequences from the Comprehensive Antibiotic Resistance Database (CARD, v.3.0.0, (10)) and the genome sequence of *T. Thermophilus* which was spiked into the samples. At the time of writing, CARD contained 2,439 AMR sequences. As CARD is primarily designed for genomic data, each sequence has an associated ‘model’ of detection i.e. criteria determining matches to the CARD reference sequences for any given query sequence. The chief distinction is between genes that have a “protein homolog” model, where detection is assessed using a BLASTP cut-off to find functional homologs (n=2,238; e.g. NDM-1 beta-lactamase), and those with a “non protein homolog” model, where detection is assessed using other methods including the locations of specific SNPs (n=247; e.g. *M. tuberculosis gyrA* conferring resistance to fluoroquinolones). Although we use a mapping-based approach from shotgun metagenomic reads, we have included this information in ResPipe. For simplicity, we designate “protein homolog” model genes and “non protein homolog” model genes under the broad headings “resistance by presence” and “resistance by variation”, respectively (where “variation” can encompass SNPs, knockout, or overexpression). The BAM files generated by the mapping were processed by a custom script to generate a count table where only alignments with a strict 100% sequence identity (without allowing any deletions or insertions) to CARD sequences were counted. Where a read mapped to more than one AMR gene family or an AMR allelic variant (i.e. could not be designated into any one AMR gene family or AMR allelic variant) it was counted as “multiple families” or “multiple alleles” respectively. For each AMR allelic variant, we calculated “lateral coverage”, defined as the proportion of the gene covered by at least a single base of mapped reads. Where reads mapped to multiple families or alleles, lateral coverage could not be calculated.

### Rarefaction curves

For fitting the relationship between sequencing depth per sample *d* and the richness *r* of AMR gene families or allelic variants, we used the species accumulation model defined by Clench (40): 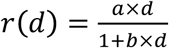. This model may be flawed, but is only used here to give a rough estimate of the sequencing depth required to achieve a proportion of *q* (e.g. 95%) of the total richness, which is then 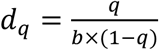.

### Normalisation of gene counts

Assuming random sequencing, longer genes are more likely to be represented in reads. In order to alleviate this gene length bias, the resulting table was adjusted by multiplying each count by the average length of mapped reads followed by dividing by the length of the AMR allelic variant to which the reads were mapped. Where there were multiple alleles, average length was used. In order to adjust for varying amounts of sample input and extraction bias, the table was further normalised to the number of reads that mapped to *T. thermophilus* using an adopted protocol from Satinsky et al. (33). We added 12.5ng of *Thermus thermophilus* to each sample. This corresponds to adding 6,025,538 copies of the *T. thermophilus* genome. The size of the *T. thermophilus* genome is 1,921,946 bases, so the number of bases of *T. thermophilus* added is 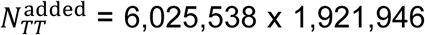. To obtain the number of bases of *T. thermophilus* recovered by sequencing 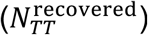, we take the number of reads assigned to *T. thermophilus* and multiply it by the insert size (300 bp). The read count *N_g_* for a particular subject *g* (e.g. a gene family or allelic variant) can then be normalised as:

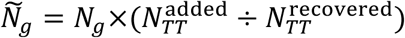

These normalisation protocols are intended to produce a pseudo-absolute gene copy number of each AMR gene family and AMR allelic variant, while recognising that this remains an estimated of the actual copy number of genes present in any given sample.

### Isolate culture and DNA extraction

For effluent samples, the effluent filter was mixed with 20 mL of nutrient broth and shaken for 10 mins at 120 rpm. 100 µL of neat sample, and 10^-1^ and 10^-2^ dilutions (in nutrient broth) were plated onto a CHROMagar Orientation agar supplemented with a 10 µg cefpodoxime disc placed on one half of the agar plate. For pig caeca and upstream sediment samples, aliquots of 100 µL of sample at neat, 10^-1^, 10^-2^, and 10^-3^-fold dilutions were plated onto a CHROMagar Orientation agar supplemented supplemented with a 10 µg cefpodoxime disc placed on one half of the agar plate. Serial dilutions were plated to enable morphological identification and isolation of individual colonies. All plates were incubated at 37 °C for 18 hours. We used cefpodoxime resistance as a surrogate marker for the selective culture of more drug-resistant *Enterobacteriaceae* (29,30).

Up to four individual colonies from each sample with a typical appearance for *Escherichia coli*, *Klebsiella* spp., *Enterobacter* spp. or *Citrobacter* spp., and from either within or external to the cefpdoxime zone, were subcultured on MacConkey agar with or without cefpodoxime discs, respectively. Following sub-culture, species was confirmed by MALDI-ToF (Bruker), and stored in nutrient broth + 10% glycerol at −80 °C prior to repeat sub-culture for DNA extraction.

DNA was extracted from pure sub-cultures using the Qiagen Genomic tip/100G (Qiagen, Venlo, Netherlands), according to the manufacturer’s instructions. Extracts from seven isolates (four from effluent, two from pig caeca, and one from upstream sediment) were selected for combination long-read (Pacific Biosciences) and short-read sequencing, based on sufficient DNA yield (with a requirement at the time of the study for ∼5µg DNA for library preparation), and appropriate fragment size distributions (assessed using TapeStation 4200, Agilent, Santa Clara, USA). These isolates were identified using MALDI-ToF as *Citrobacter freundii* (two isolates), *Enterobacter kobei/cloacae* (three isolates), and *Escherichia coli* (two isolates) (Table 1).

### Isolate sequencing

Aliquots of the same DNA extract were sequenced by two methods: short-read (Illumina), and long-read (Pacific BioSciences). For Illumina sequencing, extracts were sequenced on the HiSeq 4000 platform. Libraries were constructed using the NEBNext Ultra DNA Sample Prep Master Mix Kit (NEB), with minor modifications and a custom automated protocol on a Biomek FX (Beckman). Sequenced reads were 150 bp paired-end, with a median of 1,355,833 reads per isolate (range: 1.06-1.66 million) after read correction with SPAdes (Supplementary Table 2), corresponding to a chromosomal coverage per isolate of ∼30X with a insert size of 300 bp.

To generate long-read data from the same DNA extract for any given isolate, we used single molecule real-time sequencing using the PacBio RSII. Briefly, DNA library preparation was performed according to the manufacturer’s instructions (P5-C3 sequencing enzyme and chemistry, respectively see Supplementary Material of Sheppard et al. (31)). After read correction and trimming, there were a median of 14,189 reads per isolate (range: 12,162-17,523) with a median read length of 13,146 bp (range: 10,106-14,991) (Supplementary Table 2).

### Hybrid assembly for isolates

We assembled genomes for isolates using a version of a pipeline we had previously developed and validated against multiple *Enterobacteriaceae* genomes including two reference strains (De Maio, Shaw et al. 2019). In brief, we corrected Illumina reads with SPAdes (v3.10.1) and corrected and trimmed PacBio reads with Canu (v1.5), then performed hybrid assembly using Unicycler (v0.4.0) with Pilon (v1.22) without correction, with a minimum component size of 500 and a minimum dead end size of 500. Out of 35 total contigs across seven isolates, 28 were circularised (78%), including two chromosomes and 24 plasmids. Normalised depths of plasmids ranged from 0.6-102.6x relative to chromosomal depth, and lengths between 2.2-162.9kb (Supplementary Table 3). The majority of plasmids were found in effluent isolates (24/29). We checked MALDI-ToF species identification with mlst (v2.15.1; (32)) and found agreement (Supplementary Table 2).

### Mapping of metagenomic sequences onto isolates

To investigate the feasibility of accurately identifiying genetic structures (chromosomes and plasmids) in the metagenomic reads in relation to the impact of sequencing depth, we used the assembled chromosomes and plasmids derived from the cultured and sequenced isolates as reference genomes (*in silico* genomic “probes”) to which the metagenomic short reads were mapped. We used the same mapping protocol used for the aforementioned functional profiling and lateral coverage was calculated for each chromosome/plasmid at any given sequencing depth.

### Implementation into a Nextflow pipeline

The entire workflow (both taxonomic and AMR gene profiling) has been implemented into a Nextflow (41) pipeline complying with POSIX standards, written in Python: ResPipe (https://gitlab.com/hsgweon/ResPipe). All analyses were performed on a compute cluster hosted by the NERC Centre for Ecology and Hydrology, Wallingford, UK, with 50 compute nodes, each with a total of 1TB of RAM.

### Statistical analyses

We assessed differences in taxonomic and AMR gene profiles between replicates and sequencing lanes by calculating Bray-Curtis dissimilarities, which quantify compositional differences based on relative abundances.

These were then used to perform permutational multivariate analysis of variance tests (PERMANOVA) using the vegan package (v.2.4-1; (42)). A t-test from R base package (43) was performed to assess the differences in richness between subsampled groups of consecutive sequencing depths. Figures were produced using ggplot2 (44).

## LIST OF ABBREVIATIONS

AMR: antimicrobial resistance
SNP: single nucleotide polymorphism
CARD: (the) Comprehensive Antibiotic Resistance Database

## DECLARATIONS

### Ethics approval and consent to participate

No ethical permission was required as the samples were collected from pig caeca, at abattoir, post slaughter, by FSA or APHA vets. APHA gained permission from pig farmers/owners to obtain these samples.

### Consent for publication

Not applicable.

### Availability of data and material

The datasets generated and/or analysed during the current study are available in the NCBI repository (BioProject number: PRJNA529503). The ResPipe pipeline is available under a GPC licence at: https://gitlab.com/hsgweon/ResPipe.

### Authors’ contributions

NS and HSG designed the study, with input from all authors. DSR, MB and MA collected samples. ATH and DSR performed the laboratory work (culture, DNA extractions). RS performed PacBio long-read sequencing; Illumina sequencing was undertaken at the Wellcome Trust Centre for Human Genetics as part of a collaborative agreement. HSG developed ResPipe with input from NS; HSG and JS implemented ResPipe in NextFlow. NDM, LS, and HSG performed the data processing and bioinformatics analyses. HSG, LS, JS and NS drafted the manuscript. All authors read, improved and approved the final manuscript.

## Acknowledgements

The REHAB consortium includes (bracketed individuals included in main author list): (Abuoun M), (Anjum M), (Bailey MJ), Barker L, Brett H, (Bowes MJ), Chau K, (Crook DW), (De Maio N), Gilson D, (Gweon HS), (Hubbard ATM), (Hoosdally S), Kavanagh J, Jones H, (Peto TEA), (Read DS), (Sebra R), (Shaw LP), Sheppard AE, Smith R, Stubberfield E, (Swann J), (Walker AS), Woodford N. This publication made use of the PubMLST website (https://pubmlst.org/) developed by Keith Jolley (45) and sited at the University of Oxford. The development of that website was funded by the Wellcome Trust.

## SUPPLEMENTARY LEGENDS

**Supplementary Figure 1. Effect of sequencing depth on Bray-Curtis dissimilarity to taxonomic composition of full sample.** Results are shown for (a) Kraken and (b) Centrifuge for all samples at both genus and species level, comparing to the taxonomic composition at a depth of 200 million reads per sample.

**Supplementary Figure 2. Comparison of taxonomic classification output to simulated ground truth for CAMI gut metagenome dataset.** (a) Taxonomic composition inferred using Kraken, Centrifuge, compared to ground truth, for ten simulated samples. The top 20 most abundant genera across all samples are shown in colour. (b) Relative species abundances compared to ground truth values for Kraken (blue) and Centrifuge (red). Lines show a linear best fit.

**Supplementary Table 1. Metagenomic data.** Each sample was sequenced in replicate across four lanes (2×4=8 files per sample), combining to give the ∼200 million reads per sample used in the study. The number of reads mapping to *T. thermophilus* from each sample is also given.

**Supplementary Table 2. Hybrid sequencing details for cultured isolates.** Statistics are shown for both short reads (Illumina) and long reads (PacBio) sequenced from the same DNA extracts.

**Supplementary Table 3. Details of mapping metagenomic reads to isolate hybrid assemblies.** Each sample is shown on a different sheet.

**Supplementary Table 4. PERMANOVA results based on Bray-Curtis dissimilarities for sample replicates.** Analyses are shown in relation to sample replicates and sequencing lanes for both (a) CARD AMR abundance data, (b) Centrifuge taxonomic abundance data.

