## Supplementary figures and images for "The impact of sequencing depth on the inferred taxonomic composition and AMR gene content of metagenomic samples"

### Supplementary Figure 1

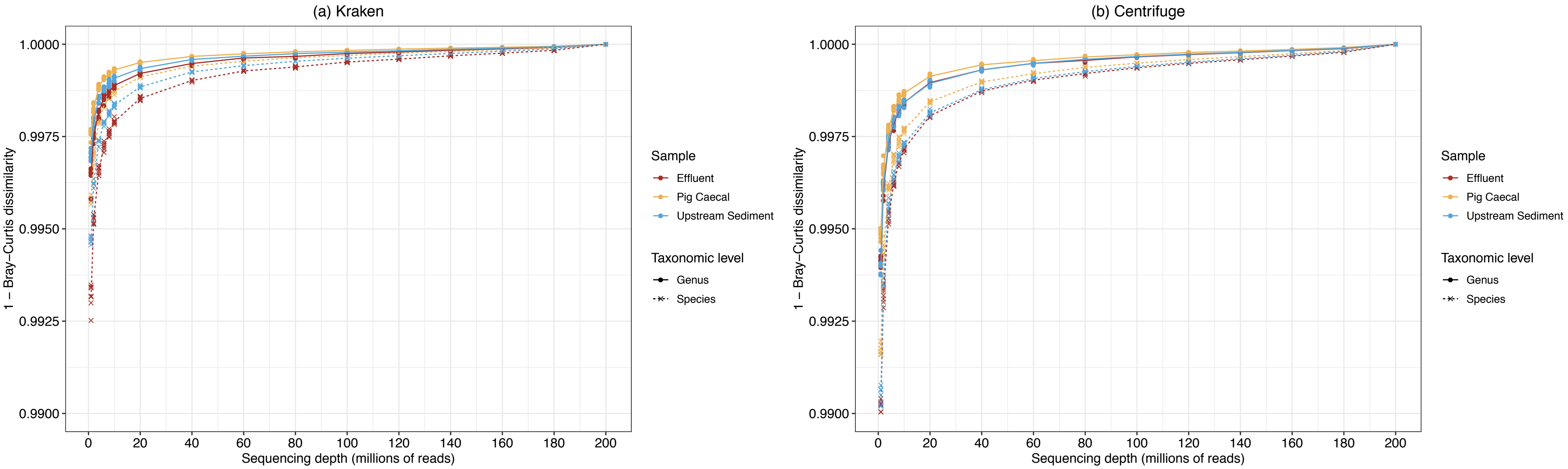
