## Supplementary Figure 2 for "The impact of sequencing depth on the inferred taxonomic composition and AMR gene content of metagenomic samples"

### (a) Taxonomic composition compared to ground truth

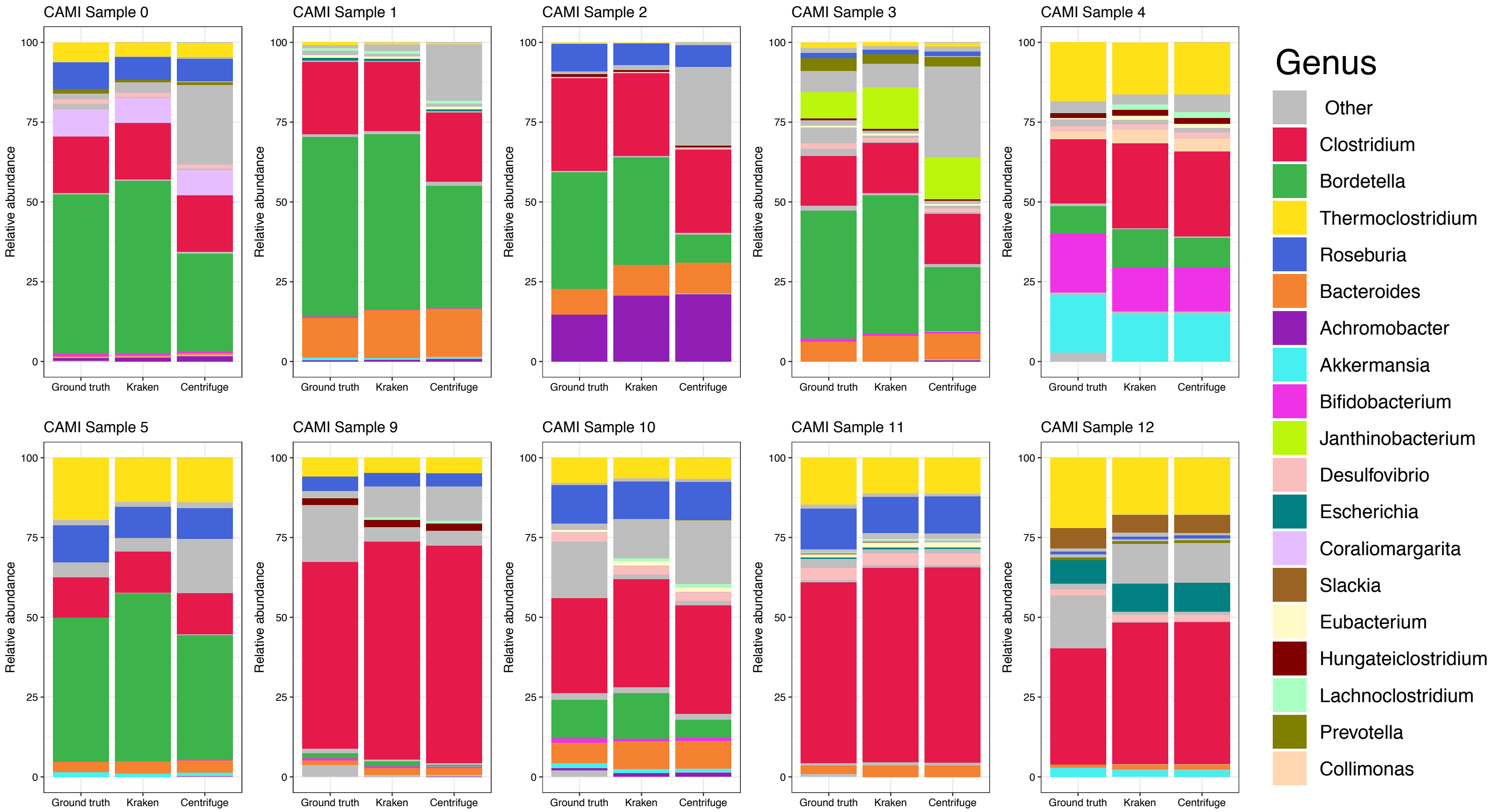


### (b) Relative species abundances compared to ground truth

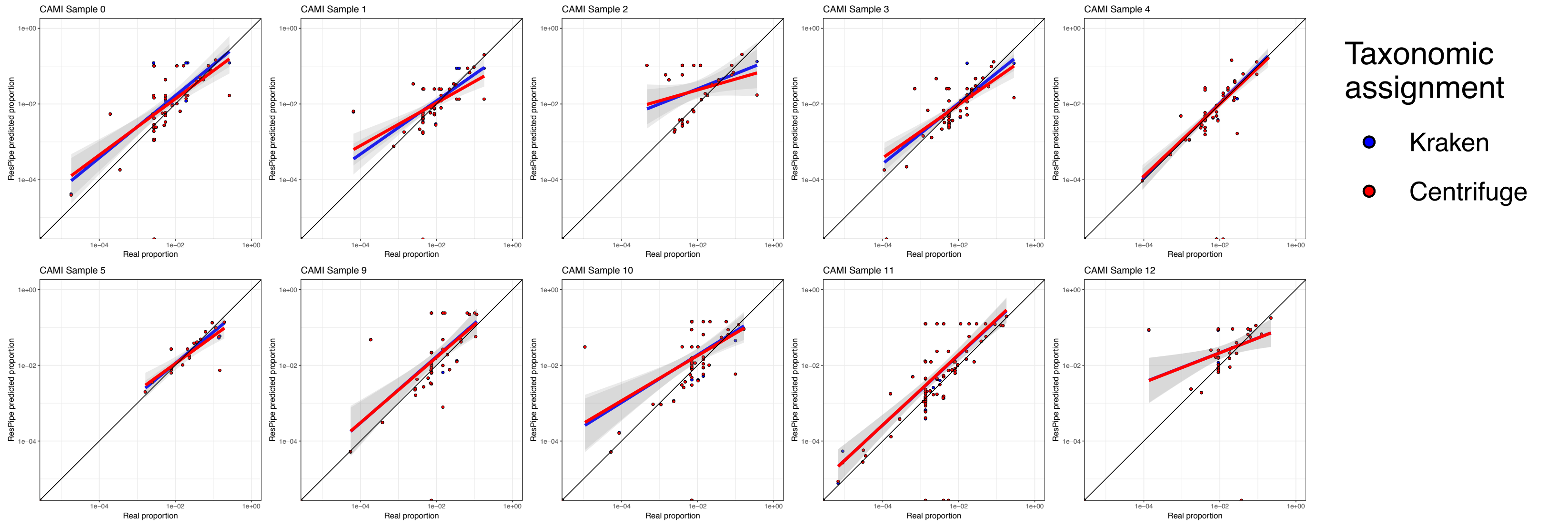
